# YeasTSS: An Integrative Web Database of Yeast Transcription Start Sites

**DOI:** 10.1101/511477

**Authors:** Jonathan McMillan, Zhaolian Lu, Judith S. Rodriguez, Tae-Hyuk Ahn, Zhenguo Lin

## Abstract

The transcription initiation landscape of eukaryotic genes is complex and highly dynamic. In eukaryotes, genes can generate multiple transcript variants that differ in 5’ boundaries due to usages of alternative transcription start sites (TSSs), and the abundance of transcript isoforms are highly variable. Due to a large number and complexity of the TSSs, it is not feasible to depict details of transcript initiation landscape of all genes using text-format genome annotation files. Therefore, it is necessary to provide data visualization of TSSs to represent quantitative TSS maps and the core promoters. In addition, the selection and activity of TSSs are influenced by various factors, such as transcription factors, chromatin remodeling, and histone modifications. Thus, integration and visualization of functional genomic data related to these features could provide a better understanding of the gene promoter architecture and regulatory mechanism of transcription initiation. Yeast species play important roles for the research and human society, yet no database provides visualization and integration of functional genomic data in yeast. Here, we generated quantitative TSS maps for twelve important yeast species, inferred their core promoters, and built a public database, YeasTSS (www.yeastss.org). YeasTSS was designed as a central portal for visualization and integration of the TSS maps, core promoters and functional genomic data related to transcription initiation in yeast. YeasTSS is expected to benefit the research community and public education for improving genome annotation, studies of promoter structure, regulated control of transcription initiation and inferring gene regulatory network.

## Introduction

Transcription initiation is the first and probably the most important step in gene expression, as it integrates the actions of key factors involved in transcription regulation (1). The short stretch of DNA immediately flanks the transcription start sites (TSSs), which is usually considered as gene’s core promoter, contain various sequence elements that accurately direct transcription initiation by the RNA polymerase II machinery (1). The accurate locations of TSSs and their transcriptional activities at a genome-scale are invaluable for many studies. For examples, they can be used for precisely determining 5’ boundary and the 5’untranslated region (5’UTR) of protein-coding genes, improving genome annotation, inference the locations of core promoters, identification of novel genes, core promoter elements, and other motifs associated with transcription (2). In addition, because most genes have multiple core promoters that have distinct activities in response to environmental cues or in different types of cells (2–5), quantitative TSS maps obtained from various growth conditions in a species are important for studying regulated control of transcription initiation and inferring gene regulatory network.

Many techniques has been applied to generate genome-wide TSS maps, such as microarray (6–8), SAGE (serial analysis of gene expression) (9), sequencing of full-length cDNA clones (10), RNA sequencing (11,12), Cap analysis gene expression (CAGE) technique (13,14), Transcript isoform sequencing (TIF-seq) (15) and Transcription start site sequencing (TSS-seq) (16). Some of these techniques, such as CAGE, TIF-seq and TSS-seq, utilize “cap-trapping” technology to pull down the 5’-complete cDNAs reversely transcribed from the captured transcripts. Through a massive parallel sequencing of the 5’ end of cDNA and analysis of the sequenced tags, the genome-wide TSSs can be identified at a single-nucleotide resolution (14). In addition, the number of mapped tags also provide a quantitative measure of transcript production from each TSS (14). CAGE is probably the most popular TSS interrogation technique and has been used to profile the locations and activities of TSSs in human (17), mouse, fruit fly and zebrafish (4,13,18) and the budding yeast *Socchoromyces cerevisiae* (2).

The high-resolution quantitative TSS maps in many organisms significantly improve our understanding of the complex and highly dynamic nature of transcription initiation landscape in eukaryotes (2–5). These studies revealed that most eukaryotic genes contain multiple core promoters which are differentially used among different tissues or developmental stages or in response to environmental cues (2,3,5,17). The selection and activity of core promoters are precisely regulated to ensure that a correct transcript is produced at an appropriate level in different tissues, developmental stages, or growth conditions. It has been shown that misregulation of transcription initiation can cause a broad range of human diseases, such as breast cancer, diabetes, kidney failure and Alzheimer’s disease (19–23). For instance, the tumor repressor gene BRCA1 has two TSSs, which produce longer and shorter transcripts. The longer BRCA1 transcript is only present in breast cancer tissues, resulted in significantly reduced production of protein products due to 10-fold less of translation efficiency than the shorter transcript (22). Unlike the translational process, which starts at the same methionine codon AUG, transcriptions are initiated from many nearby nucleotide positions in most core promoters. The transcription activities of different TSSs within a core promoter could be very different. Based on the distributions of TSS activities, the core promoters can be classified to different shape groups, which has been shown to be related to different regulatory activities (3). In addition, it was found that transcription initiation is pervasive, and it can be initiated from unconventional sites, such as exons (2,3), which generate transcripts that truncate or eliminate the predicted protein product.

Transcription initiation of eukaryotic genes is a complex and highly regulated process. Transcription of protein-coding genes is carried out by RNA polymerase II (Pol II). The recruitment of eukaryotic Pol II to a core promoter is facilitated by general transcription factors. The most-extensively studied core promoter element in eukaryotes is TATA box, which is mostly found 25-30 base pairs upstream from the TSS in mammals. However, the locations of TATA box in *S. cerevisiae* promoter was found located in a wide range of 40-120 bp upstream of TSS (24). Gene-specific transcription factors may function as activators and repressors to further increase or reduce the transcription level by binding to gene-specific transcription factor binding sites (TBFSs). TFBSs are highly enriched around 115 bp upstream of the TSS in *S. cerevisiae* (25). The presence of nucleosome in the promoter regions may be an obstacle for transcription initiation. Transcription initiation typically occurs near the boundaries of nucleosome-free regions (NFRs) (8). Several studies have found that genes with different expression profiles are associated with distinct nucleosome occupancy patterns in the promoter regions (26,27). The promoters of constantly expressed genes usually contain a nucleosome-depleted region where most TFBs are located (28,29). In contrast, conditionally expressed genes, such as stress-response genes, are associated with nucleosome-occupied promoters (27). The activation of transcription is also accompanied by alteration of chromatin structure, such as ATP-dependent nucleosome sliding (30,31) or histone modifications (32). Methylation of lysine residues in histone H3 could increase transcription by weakening chemical attractions between histone and DNA and enable the DNA to uncoil from nucleosomes (32).

To obtain genes’ TSS or 5’boundary information, most researchers rely on genome annotation files conforming to the GTF (General Transfer Format) or GFF (General Feature Format). Although a few alternative TSSs may be provided in the genome annotation files, it is not feasible for the annotation files to include the detailed information of all TSSs, such as the number and activities of TSSs within and a core promoter, regulated activities of core promoters in different cell types or under different growth conditions, and core promoter shape. Therefore, it is necessary to visualize the TSS maps to obtain a more intuitive, accurate, informative picture of transcription initiation landscape. In addition, due to the complexity of transcription initiation in eukaryotic cells, integration and visualization of functional genomic data related to transcription initiations to the TSS maps provides an intuitive illustration of the relationships between the structural components and functional elements, which could facilitate future studies about regulatory mechanism of transcription initiation. The visualization also serves as useful tools for teaching and public education to demonstrate the complexity and dynamic of transcription initiation of eukaryotic genes.

The human and mouse TSS maps can be visualized through “The FANTOM web resource” (33) and “ZENBU” (34). ZENBU also provide data integration, data analysis, and visualization system enhanced for RNA-seq, Chip-Seq, and other types of high-throughput data (34). To our knowledge, there is no web resource dedicated for visualization and integration of functional genomics data related to the transcription initiation landscape for the yeast species. Many yeast species have been served as important model organisms, widely used in daily human life, as workhorse for food, brewery, and energy industries. Particularly, the budding yeast *S. cerevisiae* has served as eukaryotic model organisms for many landmark discoveries in gene regulation mechanisms and other cellular processes over the past several decades (35). In this study, we generated high-resolution TSS maps for twelve important yeast species with divergence times ranging from 5 to 300 million years (36). We inferred core promoters for each of these species based on the quantitative TSS maps. We built a new web database: YeasTSS (www.yeastss.org), to provide free access, visualization, integration of the TSS maps, predicted core promoters, as well as other functional genomic data related to transcription initiation for yeast species. This depth and breadth of this database are valuable for the research community to improve the genome annotation of these yeast species, to study the regulated and evolutionary dynamic of transcription initiation landscape and its underlying genomic changes.

## Methods and Materials and

### Generation of TSS maps and inferences of core promoters

TSS maps and inferred core promoters are the core data of YeasTSS. We generated the TSS maps for the 12 yeast species were generated using a revised CAGE protocol, called no-amplification non-tagging CAGE libraries for lllumina sequencers (nAnT-iCAGE) (37). The nAnT-iCAGE protocol does not involve PCR amplification or restriction enzyme digestion (37), which reduced the bias created by the two processes. The cells of each species were inoculated and grown to mid-log phase in Yeast Extract-Peptone-Dextrose (YPD) Medium at 30°C for total RNA isolation. Two biological replicates of nAnT-iCAGE libraries for each species. All nAnT-iCAGE libraries were sequenced using Illumina NextSeq 500 (single-end, 75-bp reads) at the DNAFORM, Yokohama, Japan. A total number of 838,624,674 reads were generated from the 24 libraries (Table 1). The raw CAGE sequencing data have been deposited in the NCBI Sequence Read Archive (SRA) (Bioproject accession number PRJNA510689).

**Table 1.**
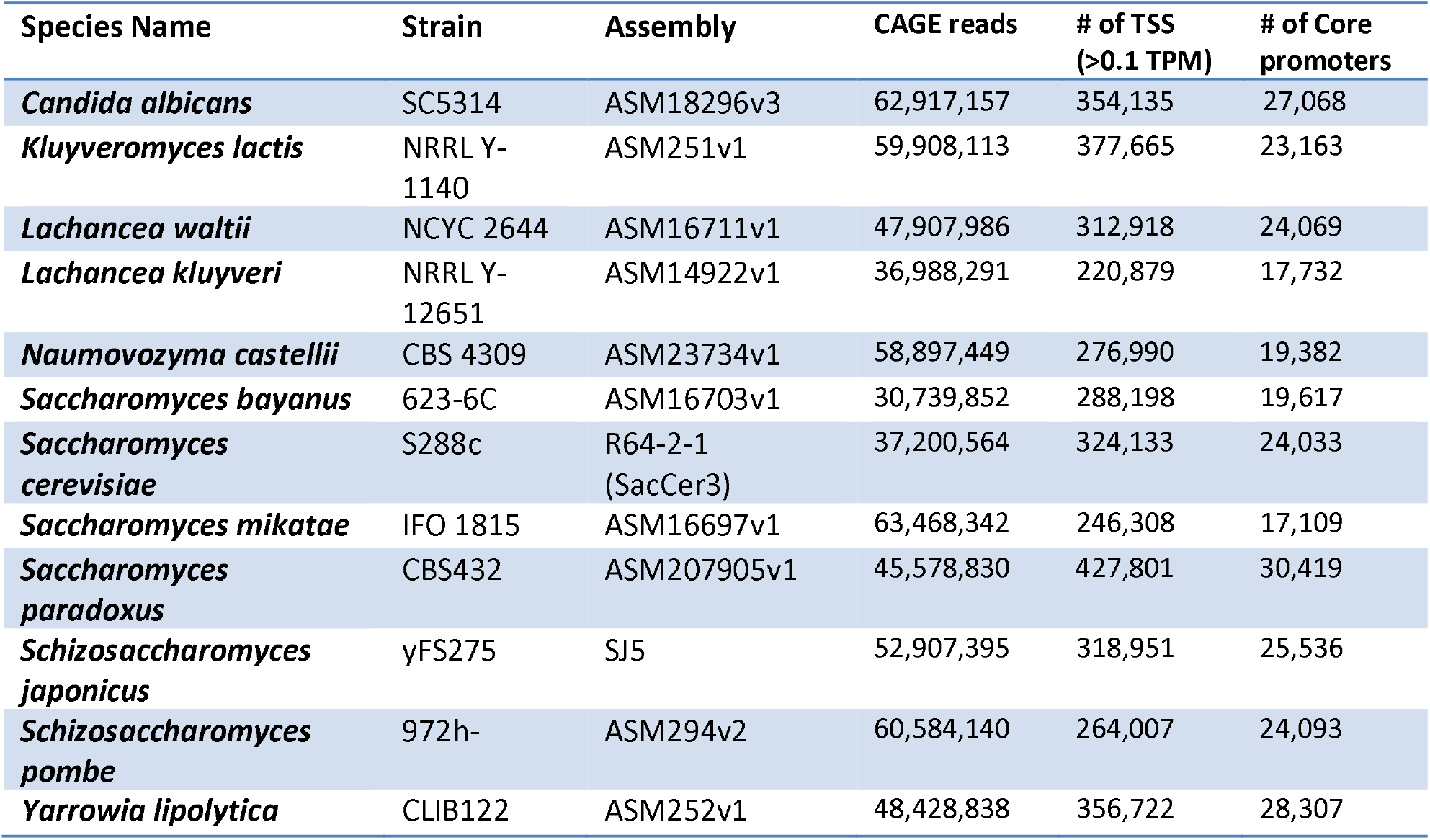
Yeast species, genome assembly and CAGE data

We identified TSS and infer core promoter based on the nAnT-iCAGE sequencing reads using a protocol described in (2) with necessary modifications. Briefly, CAGE tags were mapped to the reference genome of each species using HISAT2 (38) with no soft-clipping allowed. If the annotation of rRNA is available, CAGE tags mapped to rRNA were removed using rRNAdust (http://fantom.gsc.riken.ip/5/sstar/Protocols:rRNAdust). Only those uniquely mapped tags, (621,401,708) were used for TSS identification. CAGE-detected TSS (CTSS) were defined after correcting systematic G nucleotide addition bias at the 5’end of CAGE tags. To filter background noise, we only considered those CTSSs with tag per million (TPM) >= 0.1 for core promoter identification. A core promoter defined in this study is a cluster of nearby TSSs in a small genomic region, which reflect the transcriptional activity from the same set of general transcriptional machinery. Specifically, TSSs within 20 bp of each other were clustered as tag clusters (TCs). We then calculated the distribution of CAGE signals within a TC and used the positions of 10% quantiles and positions of 90% quantiles of CAGE signals as the boundaries of this TC. If the boundaries of two TC are < 50 bp, TCs were then aggregated together into a single set non-overlapping consensus clusters, corresponding putative core promoters.

The quantitative TSS maps and core promoters of *S. cerevisiae* in nine growth conditions were obtained from our previous study (2). These condition-specific TSS maps were generated using *S. cerevisiae* BY4741 strain (*MATa his3Δ1 leu2Δ0 met15Δ0 ura3Δ0*).

### Sources of genome sequence, annotation, and functional genomic data

The reference genome sequence and the annotation files were obtained from the *Saccharomyces* Genome Database (SGD) (39), NCBI Genome, or Yeast Gene Order Browser (YGOB) (40) (Table 1). In addition to the TSS maps generated in this study, YeasTSS also include 5’boundary of transcripts data of *S. cerevisiae* (15,16,41) and *Sch. pombe* (42) obtained from other studies. We integrated multiple functional genomics data related to transcription initiation regulation and promoter structures in several well-studied species (Table 2). The *in vivo* nucleosome occupancy data of *S. cerevisiae* and *C. albicans* were downloaded from Field et al. (43). We obtained the *in vivo* nucleosome occupancy data of *Sch. pombe* from Lantermann et al. (44). The locations of TFBS locations in *S. cerevisiae* were determined according to the motif-discovery algorithm from MacIsaac *et al.* (45), which is based on reanalysis of Chip-chip data (46), and from Venters et al. (47). The TFBS data in *Sch. pombe* was retrieved from (48). The lists of TATA box motif in *S. cerevisiae* were obtained from (49). YeasTSS also integrates the genome-wide location information of RNA polymerase II, which were obtained by ChIP-chip assays (50), DNA shape features (51), histone modifications (52,53), and RNA transcript boundaries inferred from RNA-seq data (12) in *S. cerevisiae*. The types of functional genomic data, description of data and their source used in YeasTSS are shown in Table 2.

**Table 2.**
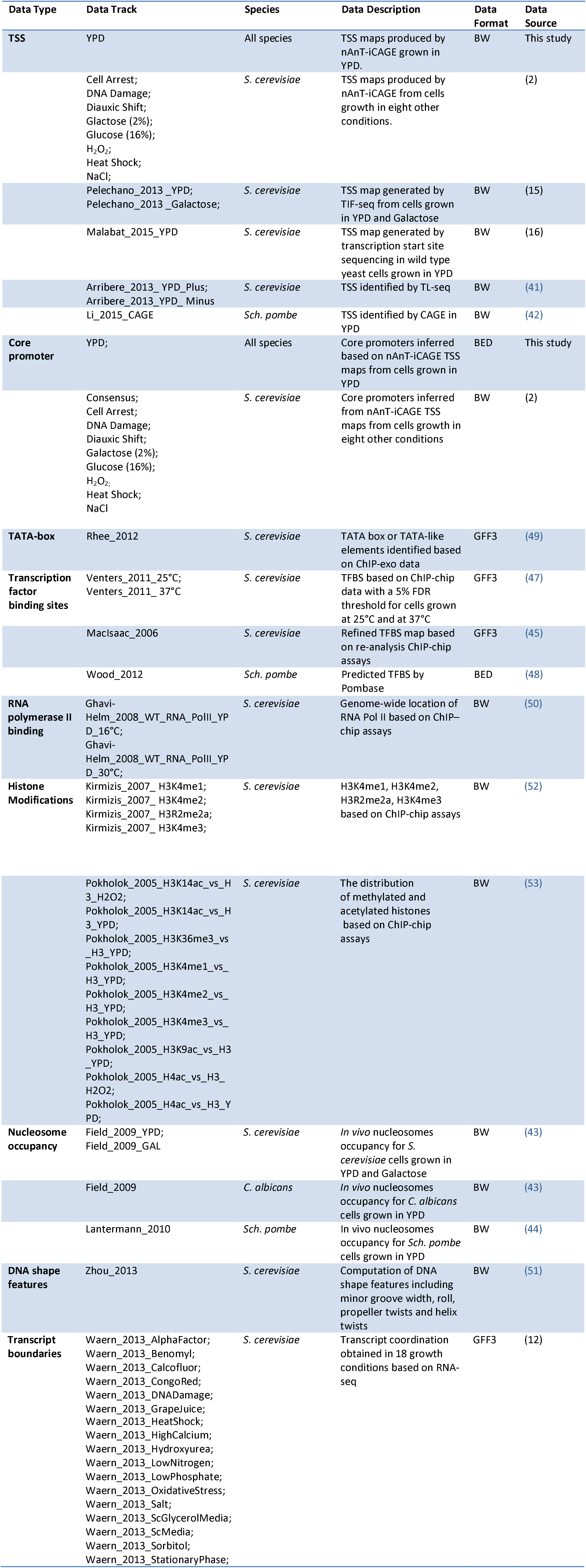
Resources and databases used for compilation and curation of data in YeasTSS.

### Data format

BED (Browser Extensible Data) format was used for functional genomic data only requires genomic coordinate information, such as the predicted core promoters, TFBS locations, TATA box, and transcript boundaries. BedGraph is a simple tab-delimited text format that is popular on most genome browser including the University of California Santa Cruz (UCSC) browser (54) for displaying the numerical value of the sequencing data at each position. However, the BedGraph text format is slow for loading and refreshing of a large data set in most genome browsers. BigWig (BW) format is a compressed binary format designed for displaying high-throughput next-generation sequencing data efficiently and effectively in genome browsers (55). We used BW format for data including both genomic coordinates and signal strength, such as quantitative TSS maps, histone modifications, and nucleosome occupancy. The “BigWig” files were converted from “BedGraph” format to allow fast and efficient remote access to the web browser server from end-users’ computers. The genomic sequence, annotation and functional genomic data in each yeast species were integrated by their genomic coordinates.

### Database development, implementation and genome browser configuration

The YeasTSS web resource was built using various computing techniques, such as Amazon Web Service (AWS), Apache web server, JBrowse utilities, PHP, Python, and Bootstrap framework. Instead of using a SQL or non-SQL database, we used various data types and in-house programs for retrieving and searching data in fast turn-around time that reduced the burden of data management and save time and costs by migrating data files to the database. The web server was built on Amazon Elastic Compute Cloud (Amazon EC2) that is an AWS web service to provide secure and resizable compute capacity in the cloud. The web server stores all data files and relevant programs for the YeasTSS database. Apache HTTP Server (Version-2.4.34) is used as a web server engine. JBrowse (Version-1.14.0), a genome browser being developed as the successor to GBrowse (56), was selected and incorporated into YeasTSS by its fast, scalable, and usable advantages. JBrowse provides fast and smooth scrolling and zooming to explore genome and sequencing information. It can scale easily to multi-gigabase genomes and deep-coverage sequencing data. The JBrowse also supports various formats of sequencing data including GFF3, BED, FASTA, Wiggle, BigWig, BAM, VCF, REST, and more. Especially, BAM, BigWig, and VCF data can be displayed directly from the compressed binary file with no conversion or extraction needed. The search functionality of the database uses PHP (Version-7.0.30) to parse the query terms and Python (Version-3.6.5) to retrieve data from the database and returns outputs in a user-friendly format. The Bootstrap (Version-4.1.1), an open source toolkit for developing with HTML, CSS, and JavaScript, is used in the YeasTSS web pages.

## Results and discussions

### YeasTSS Web interface

YeasTSS (http://www.veastss.org/) aims to provide visualization, access, and integration of quantitation TSS maps, core promoters, and various functional genomic data related to transcription initiation for important yeast species. YeasTSS provides free access and is not password-protected, and it does not require login or registration. In December 2018, the YeasTSS show twelve different yeast species (Table 1 and Fig. 1), including ten budding yeast species (such as model organism *S. cerevisiae* and human pathogen *Candida albicans*) and two fission yeast species (including another model organism *Schizosaccharomyces pombe*). The evolutionary relationships of these species, which were obtained from previous studies (36,57) and were shown by the phylogenetic tree in Figure 1. The divergence times among these species ranges from a few million years (between *S. cerevisiae* to *S. paradoxus*) to over 300 million years (between fission yeast and budding yeast) (36). The wide range of divergence times are valuable for studies of the evolutionary dynamic of transcription initiation landscape of promoter structures at different time scales

**Figure 1.**
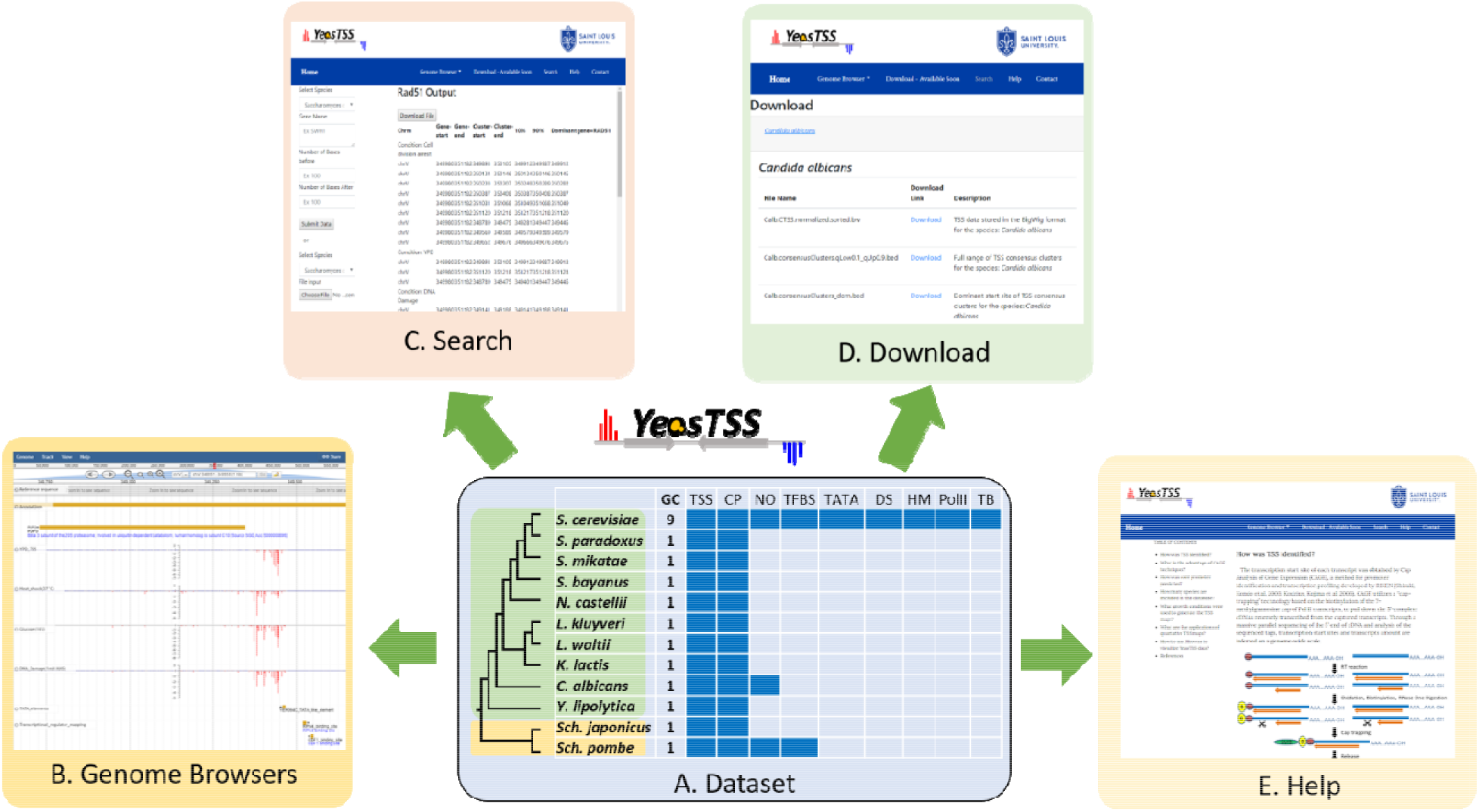
The overall design of YeasTSS. **A. Dataset:** The dataset used in YeasTSS is illustrated in this central table. Currently, YeasTSS includes twelve yeast species. The evolutionary relationships of these species are demonstrated by the phylogenetic tree on the left side of data table. The clade of 10 budding yeast species is shaded in green and the clade of 2 fission yeast species is shaded in yellow. The core promoter (CP) and TSS data of each species were generated by this study. Nucleosome occupancy (NO) data are integrated for *S. cerevisiae, Sch. pombe* and *C. albicans*. Transcription factor binding sites (TFBS) are available in *S. cerevisiae* and *Sch. pombe*. For *S. cerevisiae*, several other functional genomic data are also integrated: TATA-box, DNA shape (DS), Histone modifications (HM), Polymerase II binding (PollI), and transcript boundaries (TB) obtained from 18 different growth conditions. **B. Genome Browser:** These data are visualized and integrated by dedicated JBrowse genome browser of each species. **C. Search:** The “Search” utility provides search tools in to retrieve TSS and core promoter information from gene-by-gene analysis or global approaches. **D. Download:** The “Download” utility allows users to download all raw data used in this database through web interface. **E. Help:** The “Help” page provides documentations about of CAGE technique, TSS identification, inferences of core promoters and instructions of using genome browsers.

The homepage of YeasTSS provides a brief introduction of the database and navigation to its three major utilities: “Genome Browser”, “Search”, and “Download” (Fig. 1). These utilities can be accessed from the header bar of the homepage. The header bar is also available at all other pages of the website, so users can access and jump to any data set from any page on the website. Under the header bar of the homepage, a phylogenetic tree shows the evolutionary relationships of included yeast species. By clicking the species name in the phylogenetic tree, it will open its own dedicated genome browser page in a new window. Alternatively, the users can click the species name from the dropdown menu from “Genome Browser” in the header bar to open a new window for its genome browser. A brief description of the YeasTSS is also provided in the homepage. A detailed documentation of YeasTSS is provided by “Help” webpage, which can be accessed from the link in the header bar. The “Help” page provides information about of CAGE technique, TSS identification, inferences of core promoters and instructions of using genome browsers.

## Data visualization and integration using JBrowse genome browser

Graphical web visualization and integration of TSS maps with other functional genomic data are carried out by JBrowse genome browser in YeasTSS. Each species has a dedicated JBrowse genome browser page. Users can open a genome browser for any species in a new window by clicking the species name in the phylogenetic tree at the homepage or by clicking the species name in the dropdown menu of “Genome Browser” in the header bar. In the genome browser of each species, a user may click the chromosome names to switch the display between different chromosomes and use the left and right arrows to browse different chromosome regions. To visualize the transcription initiation landscape or promoter structure of a gene or a specific genomic region, users can enter the chromosomal coordinate ranges, gene or locus name in the search box of the genome browser. JBrowse includes a “faceted” track selector, which allows users to interactively search for the tracks they are interested in based on the metadata for each track. By clicking on the features of non-quantitative tracks, such as gene annotation, predicted core promoter, TFBS, or TATA box, users will be brought to the detail description page of the features and its associated nucleotide sequences.

To illustrate how YeasTSS can be used to visualize the complex and dynamic transcription initiation landscape and promoter structure, we demonstrate an example of using YeasTSS JBrowse genome browser to explore the *LAS17* (YOR181W) gene in *S. cerevisiae* gene (Fig. 2). *LAS17* encodes an actin-binding protein involved in actin filament organization and polymerization (58). To simplify the illustration, only a few tracks in faceted track selector were selected. As shown in Fig. 2, two predicted core promoters, which are about 500 bp part, are present in the upstream of *LAS17* ORF, suggesting that two different sets of transcription machinery may be used to initiate the transcription of *LAS17*. Here, we called the core promoter that is more proximal to the *LAS17* ORF as “LAS17-CP1”, while the distal one as “LAS17-CP2”. Transcription from the two core promoters generates transcripts isoforms with distinct lengths in the 5’ untranslated region (5’UTR). The presence of multiple core promoters in a gene has been shown to be prevalent in *S. cerevisiae* (2) as well as in mammals (3).

**Figure 2.**
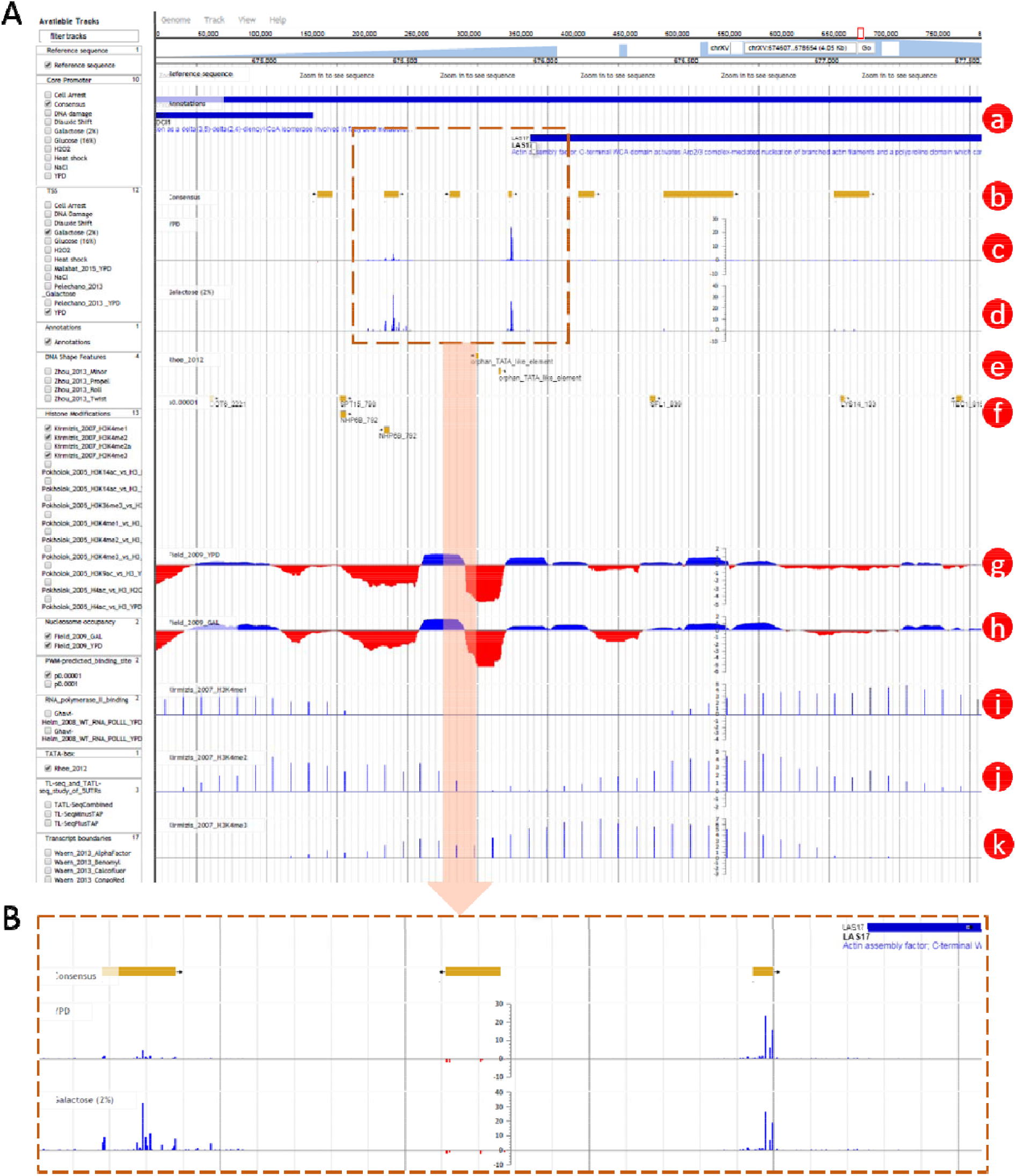
An Example of using YeasTSS Genome Browser to explore transcription initiation landscape. **A.** A 4.05 kb region around the core promoter region of LAS17 on chrXV (674,500-677,500). The available tracks in *S. cerevisiae* are provided in the left panel of genome browser. Only a few tracks were selected in this case. The tracks include: a. Genome annotation; b. Consensus clusters; c. TSS map under YPD condition; d. TSS map under YPGal (Galactose 2%) condition; e. TATA-box; f. PWM predicted binding sites; g. Nucleosome occupancy under YPD condition; h. Nucleosome occupancy under YPGal condition; i. Histone modification H3K4mel; j. Histone modification H3K4me2; k. Histone modification H3K4me3. **B.** Zoomed in region of 625 bp around the core promoter region of LAS17 on chrXV (675,375-676,000). Two consensus core promoters on the forward strand are present within 1000 upstream of LAS17. The transcription activity of each core promoter can be visualized by the TSS tracks. Different core promoter activities can be observed between the YPD and YPGal growth conditions in *S. cerevisiae*.

The detailed transcription activity within a core promoter is visualized by the TSS track, which is shown as XY-Plot wiggle track. The transcription activity from each TSS is normalized as tag per million (TPM). The positive TPM values represent transcripts generated from the forward strand, while negative TPM values indicate transcription signal from the reverse strand. The TPM values on the y-axis scale of XY-Plot wiggle track is auto-scaled to the range of TPMs in the displayed window. Within a core promoter, transcription is initiated from a cluster of nearby TSSs, rather than a single TSS, which generates transcripts with slightly different lengths. A dominant TSS that account for the majority of transcriptional activities is found in most core promoters. The distributions of TSS activity signals are very different among core promoters, forming different promoter shapes, ranging from sharp to broad. It has been found that core promoter shapes are associated with distinct regulatory patterns (3,4). In this case, LAS17-CP1 demonstrates a sharper shape than LAS17-CP2. Interestingly, the transcription activities of LAS17-CP1 are much stronger than that of LAS17-CP2 grown in rich medium (YPD). However, the relative transcription activities from the two core promoters have a significant shift if the glucose in the growth medium is replaced by 2% galactose. This phenomenon is called core promoter shift, which is found prevalent in yeast (2), mammals and fruit fly (59,60). Core promoter shift generates transcripts with distinct 5’UTRs, or results in usage of alternative translation start codon, which were speculated to have significant impacts on the translation efficiency (61,62) or mRNA stability (63). According to the in vivo nucleosome occupancy measured in yeast cells grown in YPD and galactose, both core promoters are located near the 3’end of nucleosome-depleted regions (NDR), which is consistent to the genome-wide observations (8). Different H3 methylations patterns are shown near the core promoter of LAS17. For instance, strong H3K4me1 signals are detected in the entire ORF of LAS17, while H3K4me2 signals are enriched in the 5’end of ORF. In contrast, H3K4me3 signals are mainly detected upstream of LAS17 ORF, corresponding to the promoter regions, suggesting that different histone methylations might occurs at different regions of a gene. In addition, two putative TATA-box motifs are present near 5’end of LAS17-CP1, while several TFBS motifs are found in the proximal region of LAS17-CP2, suggesting that probably different regulatory mechanisms were involved in the usage of the two core promoters.

## Search and download utilities of the YeasTSS

The “Search” utility provides search tools to retrieve TSS and core promoter information from gene-by-gene analysis or global approaches. The users have two ways to receive the core promoter and TSS data by using the search utility. First, a user can simply select a species and provide gene name(s) in the search box. A user may define the genomic range by providing specific numbers of bp before and after the start position of gene’s ORF (default values 100 bp). Alternatively, a user can provide a gene list in comma-separated values (CSV) format that should have a format of gene name, bases before gene start, and bases after gene start position. Instead of traditional SQL-based database query, this program-based search is very fast and scalable. The search output includes the genome coordinates, transcription abundance of core promoters (TPM) or tag counts of TSS within the defined searched range. The “Download” utility allows users to download all raw data used in this database through a web interface. The download page shows a brief description of each data file for users to easily capture the file content.

## Conclusions and future developments

The YeasTSS database provides a user-friendly interface for users to visualize the transcription initiation landscape and promoter structures in yeast species. YeasTSS has already been demonstrated its usefulness as a tool to support research on transcription initiation processes in yeast. YeasTSS will be periodically updated when new TSS maps or functional genomic data are available. We plan to significantly extend the capabilities of the database to increase its usefulness and functionality. Specifically, we will (i) generate TSS maps using CAGE for more yeast species, and for more *S. cerevisiae* strains, such as wild strains, wine, and pathogenic strains; (ii) integrate more functional genomics data from *S. cerevisiae* as well as other yeast species (iii) provide a graph generating tool to produce figures of any genomic region of interest for publication purpose. The data and functionality are valuable for many studies, such as analysis of the regulatory mechanism of core promote usage, the evolutionary dynamic of promoter structures, identification of 5’end boundaries of genes, improving the accuracy of genome annotation, prediction of novel genes, predictions of TFBSs, and inference of gene regulatory network. Therefore, we are confident that YeasTSS will attract a broad audience and will become a popular resource of the research community in the field of genetics, genomics and molecular biology.

## Acknowledgments

This study was supported by the start-up fund and Beaumont Award from Saint Louis University to ZL, and NSF CRII-156629, NSF-1564894, and Saint Louis University President’s Research Fund (PRF) to TA.

## References

1. Butler, J.E. and Kadonaga, J.T. (2002) The RNA polymerase II core promoter: a key component in the regulation of gene expression. Genes & development, 16, 2583-2592.

2. Lu, Z. and Lin, Z. (2018) Pervasive and Dynamic Transcription Initiation in Saccharomyces cerevisiae. bioRxiv.

3. Carninci, P., Sandelin, A., Lenhard, B., Katayama, S., Shimokawa, K., Ponjavic, J., Semple, C.A., Taylor, M.S., Engstrom, P.G., Frith, M.C. et al. (2006) Genome-wide analysis of mammalian promoter architecture and evolution. Nature genetics, 38, 626-635.

4. Hoskins, R.A., Landolin, J.M., Brown, J.B., Sandler, J.E., Takahashi, H., Lassmann, T., Yu, C., Booth, B.W., Zhang, D., Wan, K.H. et al. (2011) Genome-wide analysis of promoter architecture in Drosophila melanogaster. Genome research, 21, 182-192.

5. Raborn, R.T., Spitze, K., Brendel, V.P. and Lynch, M. (2016) Promoter Architecture and Sex-Specific Gene Expression in Daphnia pulex. Genetics, 204, 593-612.

6. Hurowitz, E.H. and Brown, P.O. (2003) Genome-wide analysis of mRNA lengths in Saccharomyces cerevisiae. Genome biology, 5, R2.

7. David, L., Huber, W., Granovskaia, M., Toedling, J., Palm, C.J., Bofkin, L., Jones, T., Davis, R.W. and Steinmetz, L.M. (2006) A high-resolution map of transcription in the yeast genome. Proceedings of the National Academy of Sciences of the United States of America, 103, 5320-5325.

8. Xu, Z., Wei, W., Gagneur, J., Perocchi, F., Clauder-Munster, S., Camblong, J., Guffanti, E., Stutz, F., Huber, W. and Steinmetz, L.M. (2009) Bidirectional promoters generate pervasive transcription in yeast. Nature, 457, 1033-1037.

9. Zhang, Z. and Dietrich, F.S. (2005) Mapping of transcription start sites in Saccharomyces cerevisiae using 5’ SAGE. Nucleic acids research, 33, 2838-2851.

10. Miura, F., Kawaguchi, N., Sese, J., Toyoda, A., Hattori, M., Morishita, S. and Ito, T. (2006) A large-scale full-length cDNA analysis to explore the budding yeast transcriptome. Proceedings of the National Academy of Sciences of the United States of America, 103, 17846-17851.

11. Nagalakshmi, U., Wang, Z., Waern, K., Shou, C., Raha, D., Gerstein, M. and Snyder, M. (2008) The transcriptional landscape of the yeast genome defined by RNA sequencing. Science (New York, N.Y, 320, 1344-1349.

12. Waern, K. and Snyder, M. (2013) Extensive transcript diversity and novel upstream open reading frame regulation in yeast. G3 (Bethesda), 3, 343-352.

13. Carninci, P., Kasukawa, T., Katayama, S., Gough, J., Frith, M.C., Maeda, N., Oyama, R., Ravasi, T., Lenhard, B., Wells, C. et al. (2005) The transcriptional landscape of the mammalian genome. Science (New York, N.Y, 309, 1559-1563.

14. Takahashi, H., Lassmann, T., Murata, M. and Carninci, P. (2012) 5’ end-centered expression profiling using cap-analysis gene expression and next-generation sequencing. Nat Protoc, 7, 542-561.

15. Pelechano, V., Wei, W. and Steinmetz, L.M. (2013) Extensive transcriptional heterogeneity revealed by isoform profiling. Nature, 497, 127-131.

16. Malabat, C., Feuerbach, F., Ma, L., Saveanu, C. and Jacquier, A. (2015) Quality control of transcription start site selection by nonsense-mediated-mRNA decay. eLife, 4.

17. Consortium, F., the, R.P., Clst, Forrest, A.R., Kawaji, H., Rehli, M., Baillie, J.K., de Hoon, M.J., Haberle, V., Lassmann, T. et al. (2014) A promoter-level mammalian expression atlas. Nature, 507, 462-470.

18. Haberle, V., Li, N., Hadzhiev, Y., Plessy, C., Previti, C., Nepal, C., Gehrig, J., Dong, X., Akalin, A., Suzuki, A.M. et al. (2014) Two independent transcription initiation codes overlap on vertebrate core promoters. Nature, 507, 381-385.

19. Arrick, B.A., Lee, A.L., Grendell, R.L. and Derynck, R. (1991) Inhibition of translation of transforming growth factor-beta 3 mRNA by its 5’ untranslated region. Molecular and cellular biology, 11, 4306-4313.

20. Romeo, D.S., Park, K., Roberts, A.B., Sporn, M.B. and Kim, S.J. (1993) An element of the transforming growth factor-beta 1 5’-untranslated region represses translation and specifically binds a cytosolic factor. Molecular endocrinology, 7, 759-766.

21. Capoulade, C., Mir, L.M., Carlier, K., Lecluse, Y., Tetaud, C., Mishal, Z. and Wiels, J. (2001) Apoptosis of tumoral and nontumoral lymphoid cells is induced by both mdm2 and p53 antisense oligodeoxynucleotides. Blood, 97, 1043-1049.

22. Sobczak, K. and Krzyzosiak, W.J. (2002) Structural determinants of BRCA1 translational regulation. The Journal of biological chemistry, 277, 17349-17358.

23. Mihailovich, M., Thermann, R., Grohovaz, F., Hentze, M.W. and Zacchetti, D. (2007) Complex translational regulation of BACE1 involves upstream AUGs and stimulatory elements within the 5’ untranslated region. Nucleic acids research, 35, 2975-2985.

24. Smale, S.T. and Kadonaga, J.T. (2003) The RNA polymerase II core promoter. Annual review of biochemistry, 72, 449-479.

25. Lin, Z., Wu, W.S., Liang, H., Woo, Y. and Li, W.H. (2010) The spatial distribution of cis regulatory elements in yeast promoters and its implications for transcriptional regulation. BMC genomics, 11, 581.

26. Jiang, C. and Pugh, B.F. (2009) Nucleosome positioning and gene regulation: advances through genomics. Nature reviews, 10, 161-172.

27. Tirosh, I. and Barkai, N. (2008) Two strategies for gene regulation by promoter nucleosomes. Genome research, 18, 1084-1091.

28. Lee, W., Tillo, D., Bray, N., Morse, R.H., Davis, R.W., Hughes, T.R. and Nislow, C. (2007) A high-resolution atlas of nucleosome occupancy in yeast. Nature genetics, 39, 1235-1244.

29. Yuan, G.C., Liu, Y.J., Dion, M.F., Slack, M.D., Wu, L.F., Altschuler, S.J. and Rando, O.J. (2005) Genome-scale identification of nucleosome positions in S. cerevisiae. Science (New York, N.Y, 309, 626-630.

30. True, J.D., Muldoon, J.J., Carver, M.N., Poorey, K., Shetty, S.J., Bekiranov, S. and Auble, D.T. (2016) The Modifier of Transcription 1 (Mot1) ATPase and Spt16 Histone Chaperone Co-regulate Transcription through Preinitiation Complex Assembly and Nucleosome Organization. The Journal of biological chemistry, 291, 15307-15319.

31. Shen, X., Mizuguchi, G., Hamiche, A. and Wu, C. (2000) A chromatin remodelling complex involved in transcription and DNA processing. Nature, 406, 541-544.

32. Shilatifard, A. (2006) Chromatin modifications by methylation and ubiquitination: implications in the regulation of gene expression. Annual review of biochemistry, 75, 243-269.

33. Kawaji, H., Severin, J., Lizio, M., Waterhouse, A., Katayama, S., Irvine, K.M., Hume, D.A., Forrest, A.R., Suzuki, H., Carninci, P. et al. (2009) The FANTOM web resource: from mammalian transcriptional landscape to its dynamic regulation. Genome biology, 10, R40.

34. Severin, J., Lizio, M., Harshbarger, J., Kawaji, H., Daub, C.O., Hayashizaki, Y., Consortium, F., Bertin, N. and Forrest, A.R. (2014) Interactive visualization and analysis of large-scale sequencing datasets using ZENBU. Nature biotechnology, 32, 217-219.

35. Duina, A.A., Miller, M.E. and Keeney, J.B. (2014) Budding yeast for budding geneticists: a primer on the Saccharomyces cerevisiae model system. Genetics, 197, 33-48.

36. Rhind, N., Chen, Z., Yassour, M., Thompson, D.A., Haas, B.J., Habib, N., Wapinski, I., Roy, S., Lin, M.F., Heiman, D.I. et al. (2011) Comparative functional genomics of the fission yeasts. Science (New York, N.Y, 332, 930-936.

37. Murata, M., Nishiyori-Sueki, H., Kojima-Ishiyama, M., Carninci, P., Hayashizaki, Y. and Itoh, M. (2014) Detecting expressed genes using CAGE. Methods Mol Biol, 1164, 67-85.

38. Kim, D., Langmead, B. and Salzberg, S.L. (2015) HISAT: a fast spliced aligner with low memory requirements. Nat Methods, 12, 357-360.

39. Cherry, J.M., Hong, E.L., Amundsen, C., Balakrishnan, R., Binkley, G., Chan, E.T., Christie, K.R., Costanzo, M.C., Dwight, S.S., Engel, S.R. et al. (2012) Saccharomyces Genome Database: the genomics resource of budding yeast. Nucleic acids research, 40, D700-705.

40. Byrne, K.P. and Wolfe, K.H. (2005) The Yeast Gene Order Browser: combining curated homology and syntenic context reveals gene fate in polyploid species. Genome research, 15, 1456-1461.

41. Arribere, J.A. and Gilbert, W.V. (2013) Roles for transcript leaders in translation and mRNA decay revealed by transcript leader sequencing. Genome research, 23, 977-987.

42. Li, H., Hou, J., Bai, L., Hu, C., Tong, P., Kang, Y., Zhao, X. and Shao, Z. (2015) Genome-wide analysis of core promoter structures in Schizosaccharomyces pombe with DeepCAGE. RNA Biol, 12, 525-537.

43. Field, Y., Fondufe-Mittendorf, Y., Moore, I.K., Mieczkowski, P., Kaplan, N., Lubling, Y., Lieb, J.D., Widom, J. and Segal, E. (2009) Gene expression divergence in yeast is coupled to evolution of DNA-encoded nucleosome organization. Nature genetics, 41, 438-445.

44. Lantermann, A.B., Straub, T., Stralfors, A., Yuan, G.C., Ekwall, K. and Korber, P. (2010) Schizosaccharomyces pombe genome-wide nucleosome mapping reveals positioning mechanisms distinct from those of Saccharomyces cerevisiae. Nature structural & molecular biology, 17, 251-257.

45. MacIsaac, K.D., Wang, T., Gordon, D.B., Gifford, D.K., Stormo, G.D. and Fraenkel, E. (2006) An improved map of conserved regulatory sites for Saccharomyces cerevisiae. BMC Bioinformatics, 7, 113.

46. Harbison, C.T., Gordon, D.B., Lee, T.I., Rinaldi, N.J., Macisaac, K.D., Danford, T.W., Hannett, N.M., Tagne, J.B., Reynolds, D.B., Yoo, J. et al. (2004) Transcriptional regulatory code of a eukaryotic genome. Nature, 431, 99-104.

47. Venters, B.J., Wachi, S., Mavrich, T.N., Andersen, B.E., Jena, P., Sinnamon, A.J., Jain, P., Rolleri, N.S., Jiang, C., Hemeryck-Walsh, C. et al. (2011) A comprehensive genomic binding map of gene and chromatin regulatory proteins in Saccharomyces. Molecular cell, 41, 480-492.

48. Wood, V., Harris, M.A., McDowall, M.D., Rutherford, K., Vaughan, B.W., Staines, D.M., Aslett, M., Lock, A., Bahler, J., Kersey, P.J. et al. (2012) PomBase: a comprehensive online resource for fission yeast. Nucleic acids research, 40, D695-699.

49. Rhee, H.S. and Pugh, B.F. (2012) Genome-wide structure and organization of eukaryotic pre-initiation complexes. Nature, 483, 295-301.

50. Ghavi-Helm, Y., Michaut, M., Acker, J., Aude, J.C., Thuriaux, P., Werner, M. and Soutourina, J. (2008) Genome-wide location analysis reveals a role of TFIIS in RNA polymerase III transcription. Genes & development, 22, 1934-1947.

51. Zhou, T., Yang, L., Lu, Y., Dror, I., Dantas Machado, A.C., Ghane, T., Di Felice, R. and Rohs, R. (2013) DNAshape: a method for the high-throughput prediction of DNA structural features on a genomic scale. Nucleic acids research, 41, W56-62.

52. Kirmizis, A., Santos-Rosa, H., Penkett, C.J., Singer, M.A., Vermeulen, M., Mann, M., Bahler, J., Green, R.D. and Kouzarides, T. (2007) Arginine methylation at histone H3R2 controls deposition of H3K4 trimethylation. Nature, 449, 928-932.

53. Pokholok, D.K., Harbison, C.T., Levine, S., Cole, M., Hannett, N.M., Lee, T.I., Bell, G.W., Walker, K., Rolfe, P.A., Herbolsheimer, E. et al. (2005) Genome-wide map of nucleosome acetylation and methylation in yeast. Cell, 122, 517-527.

54. Karolchik, D., Hinrichs, A.S., Furey, T.S., Roskin, K.M., Sugnet, C.W., Haussler, D. and Kent, W.J. (2004) The UCSC Table Browser data retrieval tool. Nucleic Acids Res, 32, D493-496.

55. Kent, W.J., Zweig, A.S., Barber, G., Hinrichs, A.S. and Karolchik, D. (2010) BigWig and BigBed: enabling browsing of large distributed datasets. Bioinformatics, 26, 2204-2207.

56. Skinner, M.E., Uzilov, A.V., Stein, L.D., Mungall, C.J. and Holmes, I.H. (2009) JBrowse: a next-generation genome browser. Genome research, 19, 1630-1638.

57. Shen, X.X., Zhou, X., Kominek, J., Kurtzman, C.P., Hittinger, C.T. and Rokas, A. (2016) Reconstructing the Backbone of the Saccharomycotina Yeast Phylogeny Using Genome-Scale Data. G3 (Bethesda), 6, 3927-3939.

58. Naqvi, S.N., Zahn, R., Mitchell, D.A., Stevenson, B.J. and Munn, A.L. (1998) The WASp homologue Las17p functions with the WIP homologue End5p/verprolin and is essential for endocytosis in yeast. Curr Biol, 8, 959-962.

59. Batut, P., Dobin, A., Plessy, C., Carninci, P. and Gingeras, T.R. (2013) High-fidelity promoter profiling reveals widespread alternative promoter usage and transposon-driven developmental gene expression. Genome research, 23, 169-180.

60. Davuluri, R.V., Suzuki, Y., Sugano, S., Plass, C. and Huang, T.H. (2008) The functional consequences of alternative promoter use in mammalian genomes. Trends Genet, 24, 167-177.

61. Kudla, G., Murray, A.W., Tollervey, D. and Plotkin, J.B. (2009) Coding-sequence determinants of gene expression in Escherichia coli. Science (New York, N.Y, 324, 255-258.

62. Livingstone, M., Atas, E., Meller, A. and Sonenberg, N. (2010) Mechanisms governing the control of mRNA translation. Phys Biol, 7, 021001.

63. Harigaya, Y. and Parker, R. (2016) Codon optimality and mRNA decay. Cell Res, 26, 1269-1270.

